# Dynamic neural representations of scene beauty are relatively unaffected by stimulus timing and task

**DOI:** 10.1101/2025.04.22.649954

**Authors:** Sanjeev Nara, Lara Becker, Lilly Hillebrand, Rongming Xiang, Daniel Kaiser

## Abstract

Understanding the neural correlates of aesthetic experiences in natural environments is a central question in neuroaesthetics. A previous EEG study (Kaiser, 2022) identified early and temporally sustained neural representations of visual scene beauty. These results were obtained with long presentation durations (1,450 ms) and with explicit beauty judgments, rendering it unclear how presentation time and task demands shape the neural correlates of scene beauty. In two EEG experiments, we replicated this study while varying presentation time and task. Experiment 1 tested whether reducing stimulus presentation time from 1,450 ms to 100 ms altered neural representations of beauty. Experiment 2 examined whether beauty-related representations prevailed when participants performed an orthogonal task instead of explicitly judging beauty. Representational similarity analysis revealed that beauty-related neural representations emerged early (within 150 – 200 ms post-stimulus) and were sustained over time, in line with previous findings. Critically, we found that neither reduced presentation time nor the absence of an explicit beauty judgment significantly altered beauty-related neural dynamics. These results suggest that the neural correlates of scene beauty are relatively robust to stimulus presentation and task regimes, providing a potential correlate of the spontaneous perception of beauty in natural environments.

## Introduction

In our daily lives, some visual inputs appear more beautiful to us than others. Understanding visual preferences for natural scenes has attracted interest in the field of neuroaesthetics. Functional magnetic resonance imaging (fMRI) studies identified a network of brain areas that represent the beauty of natural scenes, spanning the visual cortex, frontal circuits, and the default mode network (Beck and Walther, 2024; Chatterjee and Vartanian, 2014; Epstein and Kanwisher, 1998; Kawabata and Zeki, 2004; Vessel et al., 2019; Yue et al., 2007). The temporal dynamics of these representations is less well understood. A recent electroencephalography (EEG) study (Kaiser, 2022) shows that the beauty of natural scenes is represented already during early stages of cortical processing (within the first 200 ms of stimulus analysis), indicating a perceptual basis for the representation of beauty. These neural representations of beauty were temporally sustained, suggesting a continued influence of post-perceptual, cognitive processes.

This temporal cascading from prominent and early perceptual coding to sustained post-perceptual coding was established in an experimental setting where participants viewed scenes images for a relatively long duration (1,450 ms) and provided explicit beauty ratings on every trial. This raises the question how the neural dynamics related to perceived beauty change under different experimental demands. Specifically, how does the temporal evolution of representations change when presentation time and task are altered?

First, manipulations of presentation time are interesting because they limit the prolonged cognitive engagement with the stimulus while it is on the screen. In face perception, brief presentation times alter judgments of faces of low but not high attractiveness (Stróżak and Zielińska, 2019), suggesting some changes in beauty perception with shorter exposure. In scene perception, the perceived beauty of a scene is somewhat unaffected by the presentation time, with substantial correlations of beauty ratings across presentation times (Nara and Kaiser, 2024; Verhavert et al., 2018). Yet, superficially similar ratings do not exclude differences on the neural level: With shorter presentation times, beauty ratings may more strongly correspond to early neural responses reflecting perceptual processing, and less strongly to later neural responses reflecting cognitive evaluation. In Experiment 1, we test this hypothesis by replicating the previous EEG experiment on scene beauty (Kaiser, 2022) with brief presentation times of just 100 ms.

Second, manipulations of task are interesting because they probe the automaticity of beauty perception. They thereby address whether aesthetic perception is spontaneous or an intentional process (Höfel and Jacobsen, 2007). Current results on this issue are inconsistent. In face perception, EEG waveforms differentiate high and low face attractiveness, even without an attractiveness task (Halit et al., 2000; van Hooff et al., 2011). Yet, waveform differences between attractive and unattractive faces can be modulated by task (e.g., across attractiveness and gender judgments), somewhat arguing against the automaticity of beauty perception (Schacht et al., 2008). In the perception of artworks, waveform differences between artistic and non-artistic stimuli can persist under an orthogonal task regime, too (Menzel et al., 2018). In Experiment 2, we test how neural representations of scene beauty change with the current task by replicating the previous EEG experiment on scene beauty (Kaiser, 2022) with an orthogonal task unrelated to scene beauty.

In sum, we present data from two EEG experiments exploring the effect of *presentation time* and *task* on dynamic neural representations of scene beauty. Systematically comparing the results of these experiments to a previous EEG experiment with a longer presentation time and explicit beauty ratings (Kaiser, 2022), we delineate how time-resolved neural representations change with presentation time and task.

## Materials and Methods

### Participants

We conducted two EEG experiments. Experiment 1 was completed by 30 healthy adults (mean age 24.5 years, SD = 4.14 years; 21 women). One participant was excluded due to excessive motion, yielding a final sample of 29 participants. Experiment 2 was completed by 27 healthy adults (mean age 22.9 years, SD = 4.18 years; 24 women). Participants received financial remuneration. All participants provided written informed consent. Procedures were approved by the ethics committee of the Justus Liebig University Gießen and were in accordance with the Declaration of Helsinki.

### Stimuli and Paradigm

The stimulus set was the same as in our previous study (Kaiser, 2022). It consisted of 100 diverse natural scene photographs (see Fig. 1A for examples) taken from the AVA (Murray et al., 2012) and photo.net (Datta et al., 2008) image databases. The stimulus set featured 50 pairs, matched for overall content (e.g., whether animals, people, or nature was depicted) and for low-level properties (see Kaiser, 2022). The pairs were constructed so that one stimulus in each pair was judged as very beautiful in the available database ratings, whereas the other one was judged as not beautiful.

**Figure 1.**
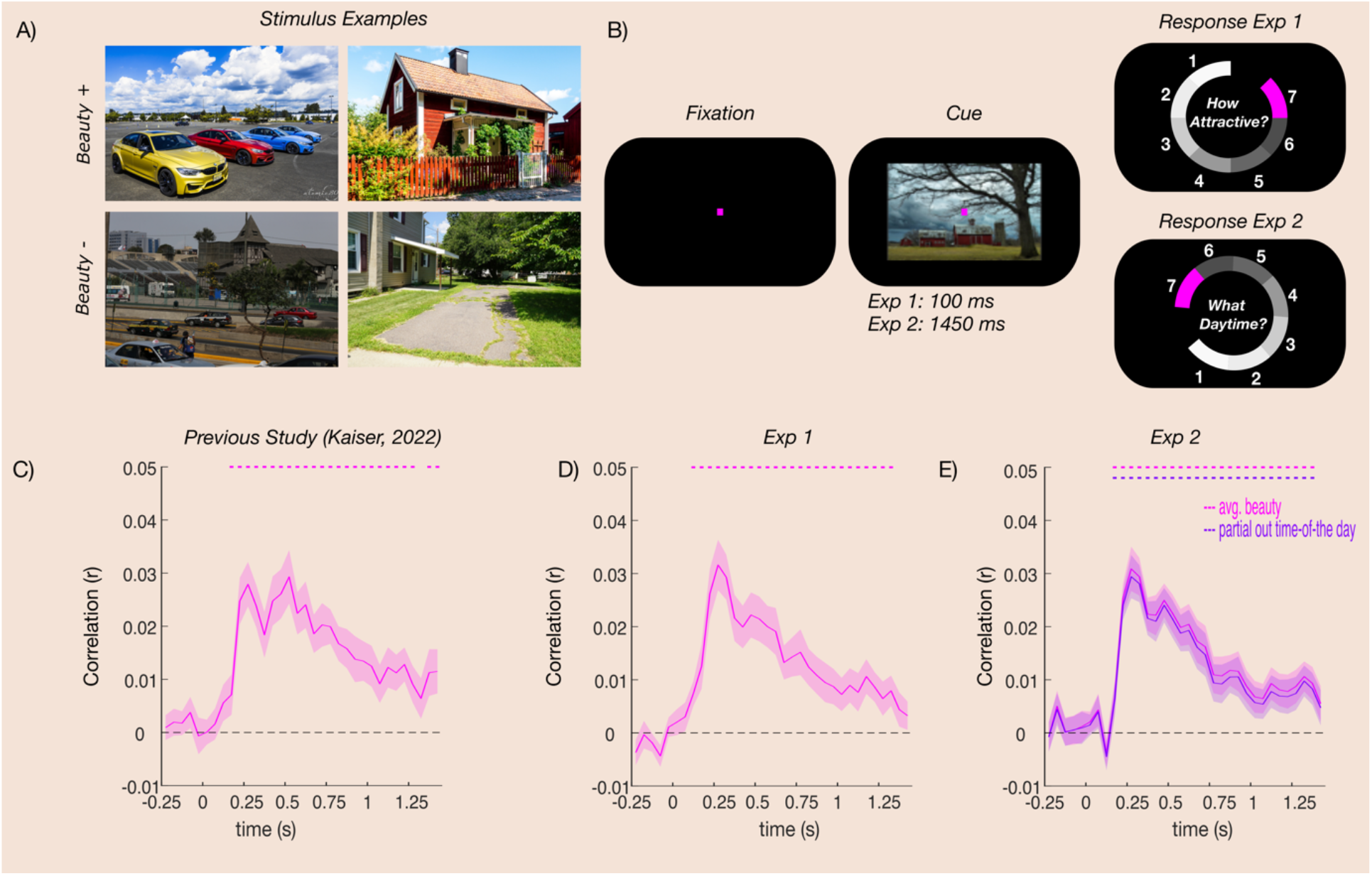
Dynamic neural representations of scene beauty across experiments. **A)** During the experiments, participants viewed 100 natural scenes depicting various contents. Scenes were arranged into 50 stimulus pairs similar in content and visual appearance, in which one stimulus was more attractive and one was less attractive. **B)** In Experiment 1, participants viewed the scenes for 100ms and provided a 1-7 beauty rating. In Experiment 2, they viewed the scenes for 1,450ms and provided a 1-7 time-of-day rating. **C)** Results from the previous Kaiser (2022) study, in which participants viewed scenes for 1,450ms and provided 1-7 beauty ratings, which we used as a benchmark to compare our results to. Here, beauty ratings predicted neural responses early on (from 100 - 150ms poststimulus) and in a sustained way. **D)** Results from Experiment 1. Beauty ratings predicted neural responses in a highly similar way as in Kaiser (2022) despite the shorter presentation regime. **E)** Results from Experiment 2. Despite the beauty-unrelated task, beauty ratings, taken from Kaiser (2022), still predicted neural responses early on and in a sustained way, even when controlling for the time-of-day ratings. Error margins reflect SEM. Significance markers denote p_corr_ < 0.05.

Both experiments used a similar paradigm, which was largely identical to the previous EEG studies on face and scene attractiveness (Kaiser, 2022; Kaiser and Nyga, 2020). On each trial, a natural scene image was presented (8° × 5.3° visual angle), followed by a 1-7 rating response screen that was operated via the computer mouse. To prevent participants from preparing a motor response in advance, the response options were presented at random angular positions across the circular response screen, with rating options (1 to 7) always increasing in clockwise direction. Participants were further instructed to keep central fixation and to restrict eye blinks to the response period. Trials were separated by an inter-trial interval randomly varying between 800 and 1200 ms. The experiment consisted of seven blocks, in each of which each scene was shown once, in random order. Each image was therefore repeated 7 times, yielding 700 trials in total.

In Experiment 1, the scene stimulus was presented for 100 ms, followed by a 1,350 ms fixation before the response. This timing was considerably faster than in the previous study (Kaiser, 2022), where the stimulus was presented for 1,450 ms, allowing us to compare beauty-related responses across conditions where sustained evaluation of the stimulus was possible or not. The stimulus was followed by a 1-7 beauty rating, as in the previous study.

In Experiment 2, the scene stimulus was presented for 1,450 ms, as in the previous study (Kaiser, 2022). Here, however, participants were asked to rate the time of day the image was taken on a scale of 1 to 7 (from 1: early morning to 7: late at night). This allowed us to investigate whether the neural correlates of scene beauty are still robustly observed without a beauty-related task.

In both studies, stimulus presentation was controlled using the Psychtoolbox (Brainard, 1997) in MATLAB 2021a.

### EEG recording and preprocessing

EEG data was recorded using a 64-channel Brain vision recorder with an Actichamp amplifier. Electrodes were placed according to the standard 10-10 system, using the Fz electrode as a reference. Data was recorded at 500 Hz. Preprocessing was performed offline using the Fieldtrip toolbox (Oostenveld et al., 2011) in MATLAB 2021a. The continuous EEG data was epoched into -500 ms to 1900 ms after the presentation of stimuli. Bad channels and bad trials were rejected through visual inspection, and eye blinks were removed using the independent component analysis (ICA) (Hyvärinen and Oja, 2000) method implemented in the Fieldtrip toolbox. The preprocessed data was then cut from -250 ms to 1,450 ms for further analyses.

### Representational Similarity Analysis

Representational similarity analysis (RSA) (Kriegeskorte et al., 2008) was used to model the EEG data. Here, we exactly replicated the RSA procedure used in Kaiser (2022) to ensure direct comparability of the results.

Neural representational dissimilarity matrices (RDMs) were created from the EEG data by computing pairwise dissimilarities for each pair of stimuli for every participant. Specifically, the EEG data were averaged within discrete time bins of 50 ms (34 bins in total), resulting in a 100× 100 RDM across 34 time points. For each time point, the data from all available time points (within a 50 ms bin) and all electrodes were used. The available data from all trials were then divided into two subsets (randomly assigned an equal number of trials per condition). The first subset was used for a PCA decomposition (to reduce the dimensionality of the response pattern). The solution of the decomposition was then projected onto the second subset, retaining components that explained 99% of the variance in the first subset. RDMs were then created from the second subset by averaging across all trials, flattening the data into response vectors for each stimulus, and computing Spearman correlation between all pairs of response vectors. The correlation values were subtracted from 1 and arranged into a 100 × 100 RDM matrix for every bin. This process was repeated again for the two subsets swapped (i.e., initially using the first subset for PCA decomposition and projecting onto the second subset, and then the reverse), and the overall procedure was repeated 50 times for each time bin, assigning trials randomly to each subset. In the end, RDMs were averaged across all 50 repetitions to yield a single RDM.

To model the neural RDMs, we created predictor RDMs from behavioral responses. For Experiment 1, predictor RDMs captured pairwise dissimilarities in beauty ratings. Here, each entry was created by computing the absolute difference between the average beauty ratings between two scenes. For Experiment 2, where participants did not rate the beauty of the scenes in the experiment, we used the average beauty rating from the Kaiser (2022) study to create the predictor RDM. We additionally created a predictor RDM based on the time-of-day ratings participants provided in the experiment, where each entry was created by computing the absolute difference in time-of-day ratings for each pair of scenes. This allowed us to control for the time-of-day ratings when assessing how beauty ratings predict the neural responses.

The correspondence between predictor RDMs and neural RDMs was assessed by correlating (Spearman correlations) all lower off-diagonal entries, separately for each time point.

### Statistical Analysis

For the RSA data, correlations between neural and predictor RDMs were compared to zero using one-sample t-tests (one-sided against zero) for each time bin. The resulting p-values were corrected for multiple comparisons using false discovery rate (FDR) corrections. Test statistics (t-values) and effect sizes (Cohen’s d) are provided for the peak effects.

For comparing the results of both Experiments to the previous results obtained with long presentation times and an explicit beauty task (Kaiser, 2022), we performed three analyses. First, we compared the correlation time courses from the RSA across studies using independent-sample t-tests at every time point and FDR-correcting the resulting p-values. Second, we compared the correlation observed at the first prominent peak of the time course (the maximum correlation value obtained between 250 – 450 ms for individual participants) across studies, using independent-sample t-tests. Finally, we fitted a linear regression model (*fitlm* function in MATLAB) to the data after the first peak to estimate how correlations were sustained across time (reflecting late perceptual as well as post-perceptual processes). The resulting linear slopes for each participant were compared across studies using independent-sample t-tests.

### Data and Code Availability

Data and code are publicly accessible on the Open Science Framework (OSF) via this link: https://osf.io/fj6sq.

## Results

### Experiment 1

To assess how beauty ratings change with presentation times, we first computed the Spearman-correlation between the average ratings obtained in Kaiser (2022) and the ratings obtained in Experiment 1. Beauty ratings were highly correlated across studies (*r* = 0.93), suggesting a limited influence of presentation time on perceived beauty (Nara and Kaiser, 2024; Verhavert et al., 2018).

To track the emergence of neural responses related to natural scene attractiveness in Experiment 1, we correlated neural RDMs with predictor RDMs that captured the images’ pairwise similarity in beauty ratings. In Kaiser (2022), where the scenes were shown for 1,450ms, the average beauty ratings predicted neural responses in a sustained fashion, starting from the 150 – 200 ms time bin (peaking at 500 – 550 ms, *t*(22) = 5.80, *p*_*corr*_ < 0.001, *d* = 1.21; Fig. 1C). In Experiment 1, with a presentation time of only 100 ms, average beauty ratings predicted neural responses in a very similar way, also starting from the 100 – 150 ms time bin (peaking at 250 – 300 ms, *t*(28) = 6.65, *p*_*corr*_ < 0.001, *d* = 1.23; Fig. 1D).

No significant differences were found when comparing the results at each time point across the two studies (all *t (50)* < 2.33, all *p*_*corr*_ > 0.76; Fig. 2A), suggesting that timing does not substantially change the neural dynamics related to perceived beauty. No differences were observed in the peak correlations either (*t(50)* = -0.90, *p* = 0.37; Fig. 2B). Fitting a linear model onto the correlation time course after the peak, we found similar negative slopes in both experiments (*t*(50) = 0.59, *p* = 0.55; Fig. 2C), indicating that representations of beauty were similarly sustained for different presentation times.

**Figure 2.**
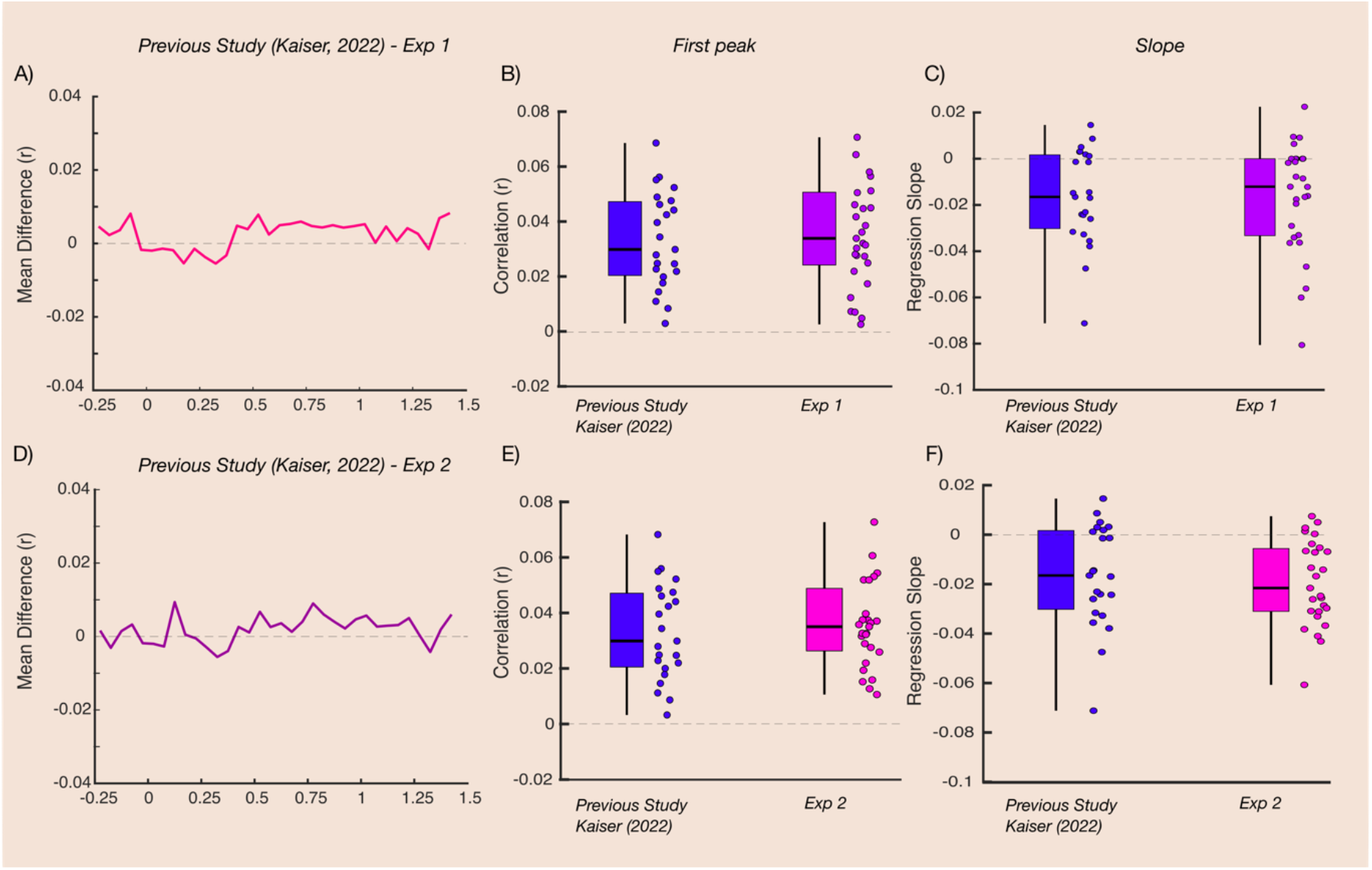
Changes in beauty-related neural representations when changing presentation time and task. **A)** Differences time course between RSA correlations obtained in the Kaiser (2022) benchmark study and Experiment 1. No significant differences were observed. **B)** RSA correlations at the first peak in the timecourse in Kaiser (2022) and Experiment 1. No significant difference was observed. **C)** Slope of a linear model fitted to the time courses after the first peak, separately for the RSA correlations in Kaiser (2022) and in Experiment 1. Again, no significant difference was observed. **D), E), F)** show the same comparisons to Kaiser (2022) for Experiment 2, again yielding no significant differences. Together, these data suggest that the neural dynamics related to perceived beauty are relatively unaffected by viewing time (Experiment 1) and task (Experiment 2).

### Experiment 2

First, to assess a possible relationship between the beauty and time-of-day ratings, we computed the Spearman correlation between the averages of the beauty ratings obtained in Kaiser (2022) and the time-of-day ratings in Experiment 3. This correlation was moderate (*r* = -0.32), suggesting that beauty ratings and time-of-day ratings were relatively independent from one another.

To track the emergence of neural responses related to natural scene attractiveness, we correlated neural RDMs with predictor RDMs that captured the images’ pairwise similarity in beauty ratings. Here, given that participants did not rate the beauty of the images, we used the average ratings for each image obtained in Kaiser (2022). Even without an explicit beauty rating task, beauty ratings predicted the neural responses from the 100 – 150 ms time bin (peaking at 250 – 300 ms, *t*(28) = 6.65, *p*_*corr*_ < 0.001, *d* = 1.23; Fig. 1E). These predictions were not driven by the time-of-day ratings across images: When partialing out similarities in time-of-day ratings, a similar prediction of the neural data from the beauty ratings was observed, again starting from the 150 – 200 ms time bin (peaking at 250 – 300 ms, *t*(26) = 7.21, *p*_*corr*_ < 0.001, *d* = 1.38; Fig. 1E). This suggests that beauty-related neural responses were not driven by a covariation in behavioral responses in the time-of-day and beauty tasks.

As for Experiment 1, no significant differences were found when comparing the results at each time point to the results in Kaiser (2022) (all *t(48)* < 2.20, all *p*_*corr*_ > 0.83; Fig. 2D), suggesting that the neural dynamics related to perceived beauty evolve in a similar way even when the beauty of the images is not task-relevant. Further, we did not find any differences in the peak correlations (*t(48)* = -0.76, *p* = 0.45; Fig 2E) or in the slope of time course after the peak (*t*(48)= 0.55, *p=* 0.58; Fig. 2F).

## Discussion

Here, we tested how dynamic neural representations of scene beauty change as a function of the stimulus presentation time (Experiment 1) and the observer’s task (Experiment 2). Comparing our results to a previous EEG study with the same scene stimuli presented with longer durations and explicitly judged for beauty, we delineated how time-varying neural representations of beauty change with presentation time and task. Strikingly, we found no substantial differences for both factors: Changing the presentation time from 1,450 ms to 100 ms, as well as changing the task from beauty judgments to time-of-day judgments, did not significantly alter the neural dynamics related to perceived beauty.

Across both experiments, we replicated the early onset of beauty-related activations within the first 200 ms of processing (Kaiser, 2022; also see Kaiser and Nyga, 2020). This early onset suggests a perceptual neural basis for evaluating beauty, in line with fMRI studies that identified neural correlates of scene beauty in the visual cortex (Nara and Kaiser, 2024; Pegors et al., 2015; Vessel et al., 2019; Yue et al., 2007). Here, we show that these perceptual representations are relatively robust to changes in presentation times, consistent with studies that report substantial correlations in ratings across stimulus durations (Nara and Kaiser, 2024; Verhavert et al., 2018). Our findings further suggest that early representations of scene beauty are not modulated by task-specific factors. They rather support the view that beauty-related visual representations emerge automatically, thereby providing a potential correlate of the spontaneous perception of beauty in real-life situations.

Perhaps more surprisingly, beauty-related representations after this initial, perceptual processing stage also seemed highly robust with respect to changes in presentation time and task. Brief presentation and an orthogonal task neither yielded less pronounced representations of beauty during later time points, nor did they alter the decay of these representations over time. The finding that the task yielded no significant alterations in beauty-related responses is somewhat difficult to reconcile with previous findings of task-specific effects in the EEG (Schacht et al., 2008), with neuroimaging work highlighting higher-order cognitive systems (Cela-Conde et al., 2004; Pegors et al., 2015; Vessel et al., 2019), as well as with theories that stress cognitive evaluation in beauty perception (Nadal and Skov, 2024). Our data at best suggest a very limited influence of task context on the neural correlates of scene beauty.

It is fair to note, however, that our stimulus set and experimental setting may have somewhat limited the potential impact of presentation time and task from the outset: First, to ensure diverse beauty ratings, we deliberately constructed our stimulus set from scenes with relatively low and relatively high aesthetical appeal: Differences in scene beauty in our experiments may thus have been easy to appreciate, even with limited exposure and without an explicit task. Second, natural scenes (in which we often spontaneously perceive beauty) may be appreciated in a more sensory-driven way than other stimulus classes like artworks (for which we appreciate beauty much more deliberately). Finally, our time-of-day task (though unrelated to beauty) perhaps was easy enough to allow participants to internally evaluate the beauty of the scenes in a deliberate manner. Perhaps more demanding tasks – in combination with brief presentation regimes – are needed to observe pronounced differences at the neural level.

When reconciling our results with previous fMRI studies, the absence of task-specific effects may also relate to a relative insensitivity of EEG signals with respect to the finer-grained representations emerging at higher levels of cortical hierarchies. Notable divergences between M/EEG and fMRI results have been reported when investigating conceptual representations in the visual hierarchy (Iamshchinina et al., 2021; Proklova et al., 2019).

Taken together, our findings indicate that dynamic neural representations of scene beauty are relatively unaffected by presentation time and task. This suggests that perceptual stimulus attributes play a major role in beauty perception for natural scenes. This dominance of perceptual attributes may relate to our tendency to spontaneously appreciate beauty in natural environments. However, more work is needed to fully understand how this robustness to presentation and task regimes generalizes across stimuli and context.

## Acknowledgements

This project was supported by the German Research Foundation (DFG; KA4683/6-1, project number 536053998) and by “the Adaptive Mind”, funded by the Excellence Program of the Hessian Ministry of Higher Education, Science, Research and Art. The authors declare that there are no conflicts of interest.

## Author contributions

Conceptualization: SN and DK

Data curation: SN

Formal analysis: SN, LB, LH and RX

Funding acquisition: DK

Investigation: SN, LB, LH and RX

Methodology: SN and DK

Project administration: SN and DK

Resources: SN and DK

Software: SN

Supervision: SN and DK

Validation: SN and DK

Visualization: SN

Writing—original draft: SN

Writing—review and editing: SN, LB, LH, RX and DK

## Competing interests

The authors declare that they have no competing interests.

## Notes

### Competing Interest Statement

The authors have declared no competing interest.

## References

Beck, D., Walther, D.B., 2024. The Natural Scene Network. Oxford Research Encyclopedia of Neuroscience. 10.1093/ACREFORE/9780190264086.013.396

Brainard, D.H., 1997. The Psychophysics Toolbox. Spat Vis 10, 433–436. 10.1163/156856897X00357

Cela-Conde, C.J., Marty, G., Maestú, F., Ortiz, T., Munar, E., Fernández, A., Roca, M., Rosselló, J., Quesney, F., 2004. Activation of the prefrontal cortex in the human visual aesthetic perception. Proc Natl Acad Sci U S A 101, 6321–6325. 10.1073/PNAS.0401427101

Chatterjee, A., Vartanian, O., 2014. Neuroaesthetics. Trends Cogn Sci 18, 370–375. 10.1016/J.TICS.2014.03.003

Datta, R., Li, J., Wang, J.Z., 2008. Algorithmic inferencing of aesthetics and emotion in natural images: An exposition, in: Proceedings - International Conference on Image Processing, ICIP. pp. 105–108. 10.1109/ICIP.2008.4711702

Epstein, R., Kanwisher, N., 1998. A cortical representation of the local visual environment. Nature 1998 392:6676 392, 598–601. 10.1038/33402

Halit, H., De Haan, M., Johnson, M.H., 2000. Modulation of event-related potentials by prototypical and atypical faces. Neuroreport 11, 1871–1875. 10.1097/00001756-200006260-00014

Höfel, L., Jacobsen, T., 2007. Electrophysiological indices of processing aesthetics: Spontaneous or intentional processes? International Journal of Psychophysiology 65, 20–31. 10.1016/j.ijpsycho.2007.02.007

Hyvärinen, A., Oja, E., 2000. Independent component analysis: algorithms and applications. Neural Networks 13, 411–430. 10.1016/S0893-6080(00)00026-5

Iamshchinina, P., Kaiser, D., Yakupov, R., Haenelt, D., Sciarra, A., Mattern, H., Luesebrink, F., Duezel, E., Speck, O., Weiskopf, N., Cichy, R.M., 2021. Perceived and mentally rotated contents are differentially represented in cortical depth of V1. Communications Biology 2021 4:1 4, 1–8. 10.1038/s42003-021-02582-4

Kaiser, D., 2022. Characterizing Dynamic Neural Representations of Scene Attractiveness. J Cogn Neurosci 34, 1–10. 10.1162/jocn_a_01891

Kaiser, D., Nyga, K., 2020. Tracking cortical representations of facial attractiveness using time-resolved representational similarity analysis. Sci Rep 10. 10.1038/S41598-020-74009-9

Kawabata, H., Zeki, S., 2004. Neural correlates of beauty. J Neurophysiol 91, 1699–1705. 10.1152/JN.00696.2003

Kriegeskorte, N., Mur, M., Bandettini, P., 2008. Representational similarity analysis - connecting the branches of systems neuroscience. Front Syst Neurosci 2. 10.3389/neuro.06.004.2008

Menzel, C., Kovács, G., Amado, C., Hayn-Leichsenring, G.U., Redies, C., 2018. Visual mismatch negativity indicates automatic, task-independent detection of artistic image composition in abstract artworks. Biol Psychol 136, 76–86. 10.1016/J.BIOPSYCHO.2018.05.005

Murray, N., Marchesotti, L., Perronnin, F., 2012. AVA: A large-scale database for aesthetic visual analysis. Proceedings of the IEEE Computer Society Conference on Computer Vision and Pattern Recognition 2408–2415. 10.1109/CVPR.2012.6247954

Nadal, M., Skov, M., 2024. The sensory valuation account of aesthetic experience. Nature Reviews Psychology 2024 4:1 4, 49–63. 10.1038/s44159-024-00385-y

Nara, S., Kaiser, D., 2024. Integrative processing in artificial and biological vision predicts the perceived beauty of natural images. Sci Adv 10. 10.1126/sciadv.adi9294

Oostenveld, R., Fries, P., Maris, E., Schoffelen, J.M., 2011. FieldTrip: Open source software for advanced analysis of MEG, EEG, and invasive electrophysiological data. Comput Intell Neurosci 2011. 10.1155/2011/156869

Pegors, T.K., Mattar, M.G., Bryan, P.B., Epstein, R.A., 2015. Simultaneous perceptual and response biases on sequential face attractiveness judgments. J Exp Psychol Gen 144, 664–673. 10.1037/XGE0000069

Proklova, D., Kaiser, D., Peelen, M. V., 2019. MEG sensor patterns reflect perceptual but not categorical similarity of animate and inanimate objects. Neuroimage 193, 167–177. 10.1016/J.NEUROIMAGE.2019.03.028

Schacht, A., Werheid, K., Sommer, W., 2008. The appraisal of facial beauty is rapid but not mandatory. Cogn Affect Behav Neurosci 8, 132–142. 10.3758/CABN.8.2.132/METRICS

Stróżak, P., Zielińska, M., 2019. Different processes in attractiveness assessments for unattractive and highly attractive faces—The role of presentation duration and rotation. Acta Psychol (Amst) 200. 10.1016/j.actpsy.2019.102946

van Hooff, J.C., Crawford, H., van Vugt, M., 2011. The wandering mind of men: ERP evidence for gender differences in attention bias towards attractive opposite sex faces. Soc Cogn Affect Neurosci 6, 477–485. 10.1093/SCAN/NSQ066

Verhavert, S., Wagemans, J., Augustin, M.D., 2018. Beauty in the blink of an eye: The time course of aesthetic experiences. British Journal of Psychology 109, 63–84. 10.1111/BJOP.12258

Vessel, E.A., Isik, A.I., Belfi, A.M., Stahl, J.L., Gabrielle Starr, G., 2019. The default-mode network represents aesthetic appeal that generalizes across visual domains. Proc Natl Acad Sci U S A 116, 19155–19164. 10.1073/PNAS.1902650116/SUPPL_FILE/PNAS.1902650116.SAPP.PDF

Yue, X., Vessel, E.A., Biederman, I., 2007. The neural basis of scene preferences. Neuroreport 18, 525–529. 10.1097/WNR.0b013e328091c1f9

